# Influenza A virus exploits the motility of membrane cytoskeletal actomyosin filaments for its genome packaging in the host cell

**DOI:** 10.1101/2024.09.11.612468

**Authors:** I-Hsuan Wang, Jiro Usukura, Yasuyuki Miyake, Eiji Usukura, Akihiro Narita, Yohei Yamauchi, Yoshihiro Kawaoka

## Abstract

Influenza A virus encodes its genome in eight segments of viral ribonucleoproteins (vRNPs) replicated in the host cell nucleus. Our understanding of host factors involved in driving vRNP selective packaging remains incomplete. To address this, we used advanced immuno-freeze-etch electron microscopy to visualise the vRNP packaging process and atomic force live-cell imaging (AFM) to examine the motility of membrane cytoskeletal actin filaments. In the cytoplasm, vRNPs were mainly localised on mottled membrane-like structures, suggesting intracellular trafficking through such structures. After reaching the cytoplasmic side surface of the plasma membrane, vRNPs formed many aggregates while associating with actin filaments. Antibody labelling also detected myosin along actin filaments entangled in vRNPs. Blocking myosin activity with blebbistatin prevented the active movement of membrane cytoskeletal actin filaments just below the plasma membrane visualised by AFM and abrogated proper aggregation of vRNPs. Thus, actomyosin motility appears to be crucial for the selective packaging of vRNPs.

## Introduction

Influenza A virus (IAV), a member of the *Orthomyxoviridae* family, poses substantial threats to both public and veterinary health^1^. Its genome is segmented, hence, facilitates the emergence of recombinant strains with pandemic potential^1–4^. The segmented genome consists of eight viral ribonucleoproteins (vRNPs), each comprising of a different viral RNA (vRNA) enveloped by a nucleoprotein (NP), the ends of which are anchored by a unit of RNA-dependent RNA polymerase (RdRp) ^5,6^. The RdRp complex consists of the polymerase acidic (PA) and polymerase basic 1 and 2 (PB1 and PB2) proteins. Each vRNA segment encodes specific viral protein(s) essential for viral replication and assembly. Additionally, the error-prone nature of IAV RNA polymerase contributes to genomic drift, resulting in the development of mutant variants capable of evading existing population immunity^7^. Consequently, these mutations play a pivotal role in initiating annual epidemics.

IAV infection in cells initiates with binding of virions to sialic acid (receptor) of glycoproteins and glycolipids in the plasma membrane. Virions attached to the plasma membrane induces receptor-mediated endocytosis and macro-pinocytosis^8,9^, which are internalized into the cytoplasm as endosomes. Endocytic vesicles (early endosomes) containing the IAV virions migrate towards the nuclear periphery as they mature into late endosome. Acidification within the late endosomes causes haemagglutinin (HA)-mediated fusion with the endosomal membrane, leading to disruption of the matrix protein (M1) layer; this process facilitates the release of vRNPs into the cytoplasm, which are subsequently imported into the nucleus via the importin-α/β2 pathway ^4,10–13^. Once in the nucleus, vRNPs use RNA-dependent RNA polymerase (RdRp) to replicate the genomic viral RNA and transcribe the viral mRNA for viral protein synthesis ^4^. Progeny vRNPs form a complex with M1 and nuclear export protein (NEP/NS2), which links them to CRM1, facilitating vRNP cytoplasmic export^4,14,15^. In the cytoplasm, vRNPs interact with Rab11 recycling endosomes to form liquid organelles known as viral inclusions near endoplasmic reticulum (ER) exit sites ^16–20^. Such viral inclusion bodies are therefore hypothesized to be sites that facilitate inter-segmental vRNP-vRNP interactions to assemble specific genomes. In a specific assembly mechanism that is not yet fully understood, progeny vRNPs are said to be ultimately transported to the budding zone on the plasma membrane via Rab 11-dependent ER for subsequent virion assembly^17–21^. Biochemical affinity between the nucleoprotein (NP) and actin filament was reported ^22,23^, suggesting the involvement of actin filaments in the IAV infection cycle. This has led to further research using immunocytochemistry and electron microscopy focusing on the role of actin filaments in vRNP transport and packaging^24–27^. However, it is still insufficient with these results to elucidate where and how vRNPs interact with actin filaments. The actual binding of actin filaments to vRNPs and their packaging processes have not yet been visualised. This study aims to reveal the actin filament-mediated packaging process of genomic vRNPs by morphological approaches using immuno-freeze-etching electron microscopy (immuno-FE) combined with unroofing techniques and live cell imaging with high-speed atomic force microscopy (AFM).

## Results

### Localization of vRNPs in the cytoplasm

Immuno-freeze etching electron microscopy (Immuno-FE) of cryo-sections was performed to determine the localization of vRNPs in the cytoplasm of A549 cultured cells at 16 hours post infection (hpi). Cryo-sections of infected cells were prepared according to the Tokuyasu method^28,29^, and labelled with anti-vRNP polyclonal antibody at room temperature. Then the sections were quick-frozen again and placed in a freeze-etching apparatus. The surface of the frozen sections was slightly freeze-dried (freeze-etched) and the structures thereby exposed were replicated by evaporation of platinum and carbon^30^ (see Methods for details). Anti-vRNP antibody binding sites (i.e., vRNPs) were predominantly localised on the surface of the characteristic membrane-like structures (Fig. 1a), where no aggregations of vRNPs was observed comparing to the cytoplasmic side of the plasma membrane as described below. These membrane-like structures were scattered in a mottled pattern throughout the cytoplasm and appeared to be diverse in size and shape when viewed under electron microscope (EM) (Fig. 1a). Some of them were more than 1 µm long and sometimes nipped in places. Interestingly, when enlarged, the membrane-like structure appeared to contain fibrous components (Fig.1b). They were also placed on a thin filament weave (Fig. 1a, b). The details of the membrane-like structure are currently unknown, including whether this is a morphological counterpart of the so-called liquid organelle^16^.

**Fig. 1.**
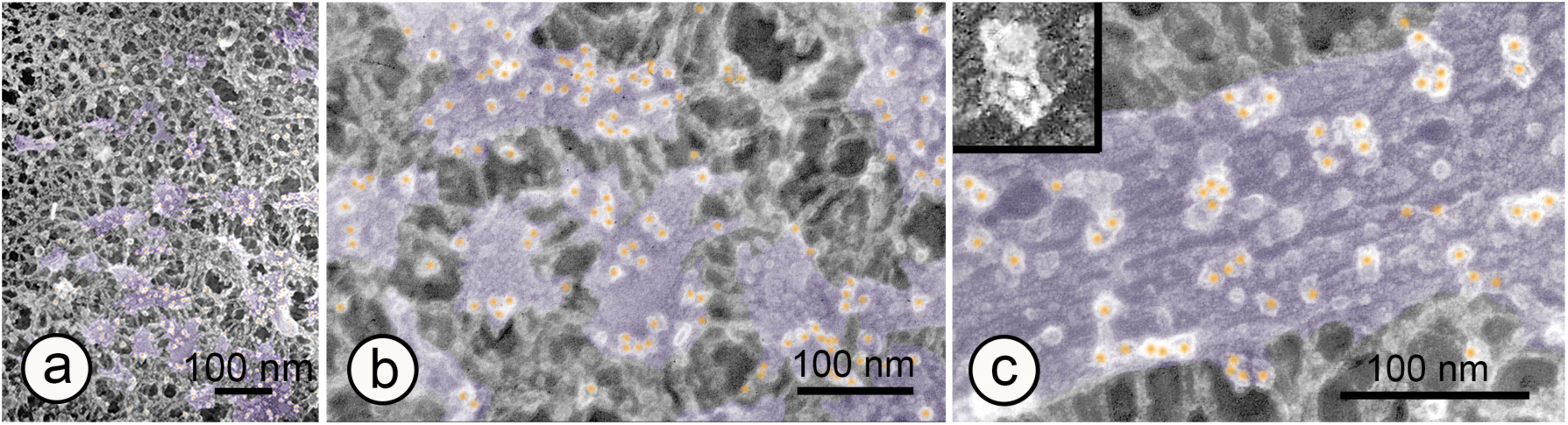
Cryo-section-immuno-FE electron micrographs of cells infected with IAV. The colloidal gold-conjugated secondary antibody is highlighted in orange. **a:** Low magnification image of immuno-FE in cryosections of cytoplasm. Membrane-like structures (overlaid with transparent violet) are patchily distributed in the cytoplasm. **b:** Anti-RNP antibody binding sites (vRNPs) are localized predominantly on the membrane-like structures (overlaid with transparent violet), that appear to be laid on the weave of thin filaments. **c:** High magnification electron micrograph of a membrane-like structure, which appears to contain fibrous components. Inset shows the colloidal gold-conjugated secondary antibody before colouring. The white sphere in the centre is colloidal gold (10 nm in diameter, shown here in negative contrast for clarity) and the irregular circles surrounding it are immuno-gamma globulins (IgG) bound to colloidal gold, which are visualized by rotary shadowing of platinum.

### Morphological approach to the packaging process of vRNPs

Morphological observations of the cortical regions of the infected cells, especially the cytoplasmic side surface of the plasma membrane, are essential to elucidate where progeny vRNPs are transported and how they are packaged into the core of the progeny virion there. Immuno-FE combined with the unroofing method^31,32^ was used for this purpose (see Methods for details). Numerous vRNP immunogold labels (i.e. vRNPs) were accumulated on the cytoplasmic side surface of the plasma membrane, forming numerous aggregates of various sizes and shapes (Fig. 2). When zoomed in, the thin filaments interconnecting the immunolabelled vRNPs became visible (Fig. 2b). These filaments were identified as actin filaments based on anti-actin antibody labelling (Fig. 3a) and their structural features (i.e. thickness of about 7 nm and periodic striations of about 5.5 nm). Since many of these actin filaments were in contact with the plasma membrane, they appeared to be part of the membrane cytoskeleton. Myosin was also detected along actin filaments by anti-myosin antibody labelling (Fig. 3b), suggesting myosin-induced movement of actin filaments. While biochemical affinity between the nucleoprotein (NP) and actin had already been reported^22,23^, the actin filaments linking the vRNPs were captured as images for the first time in this study. This is an important finding in elucidating the mechanism and process of vRNPs packaging. On thick actin bundles, antibody-labelled vRNPs tended to line up linearly, whereas vRNPs forming aggregates of various shapes were associated with thin actin filaments (Fig. 4 a-e). The thin actin filaments were often sharply curved and intricately entangled with vRNPs in the aggregate (Figs. 3,4). As shown in Figure 2b, the aggregates were also linked to each other by actin filaments (inter-aggregate actin filaments). When the vRNPs were packed tightly in the circular pattern, the inter-aggregate actin filaments collapsed, and each aggregate became independent (Fig. 2c and Fig. 4e). However, some thin, short actin filaments remained within the aggregate and intertwined with the vRNPs (Fig.4e). In fact, Western blotting also detected actin, myosin and cofilin in purified IAV (Extended data Fig. 1) as reported previously ^33, 34^. These results suggest that actin, myosin, and related proteins were used in the packaging of vRNPs and that some of them were incorporated into progeny virions while being entangled with vRNPs.

**Fig. 2.**
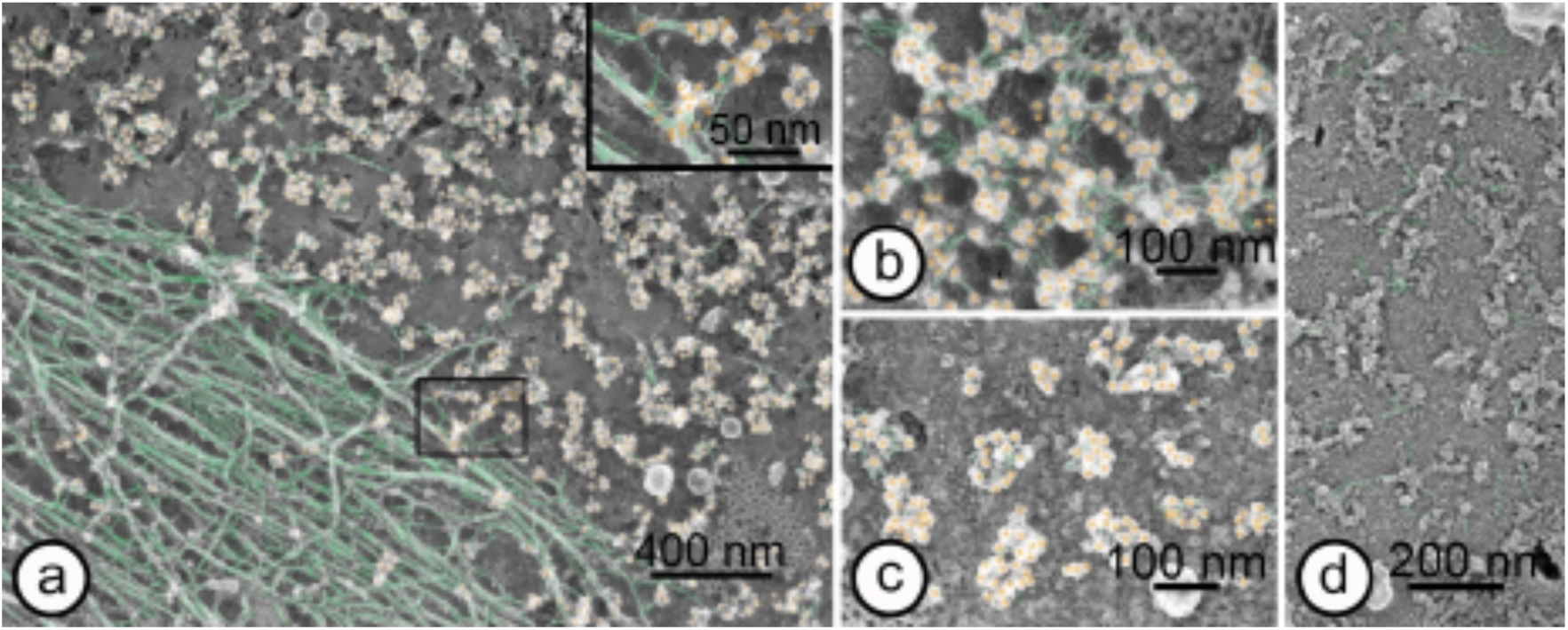
Unroofing-immuno-FE electron micrographs of cells infected with IAV. **a**: Anti-vRNP antibody labels are distributed on the cytoplasmic side surface of the plasma membrane, forming aggregates of various sizes. Very little antibody labelling (i.e., vRNPs) is observed in stress fibres, but linear arrays of vRNPs are found on the lateral twigs that are in contact with the plasma membrane (inset). Inset show enlargement of framed area. **b**: Slightly magnified immuno-FE micrograph showing actin filaments associated with vRNPs. **c**: Immuno-FE micrographs showing the labelled vRNPs assembled in circular pattern. **d**: Electron micrograph showing immuno-FE control using non-immune serum instead of primary antibody. Immuno-gold particles bound to antibodies are coloured in orange. Actin filaments are coloured in green.

**Fig. 3.**
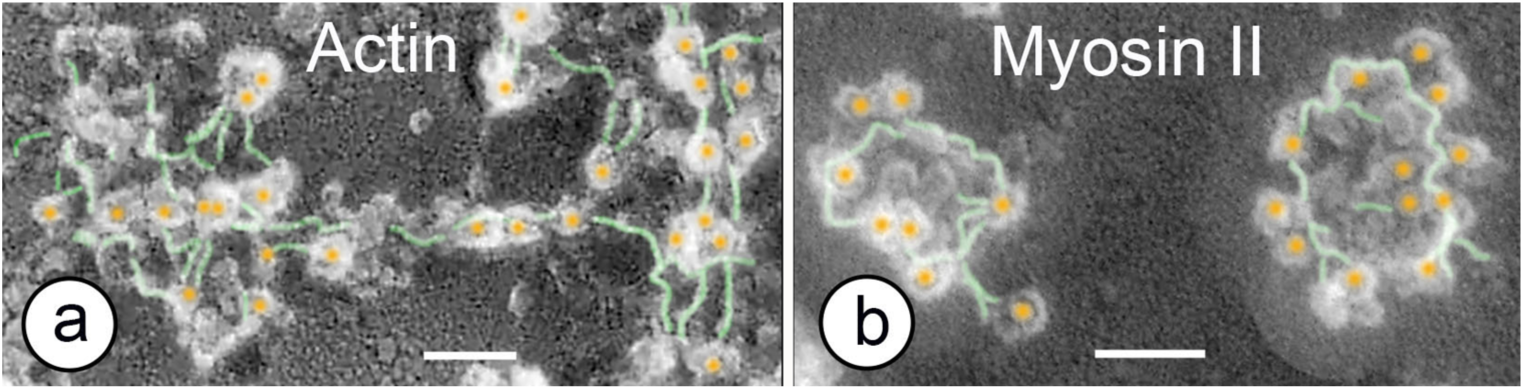
High-magnification immuno-FE electron micrographs showing immuno-localization of actin (**a**), myosin(**b**). **a**: Anti-cytoskeletal γ actin antibody labelling is also found within vRNP clusters in addition to stress fibres. **b**: Anti-myosin II antibody labels are found along actin filaments within vRNP aggregates. Colloidal gold-conjugated secondary antibodies are highlighted in orange colour. Actin filaments are coloured in green. Scale bars: 50 nm

**Fig. 4.**
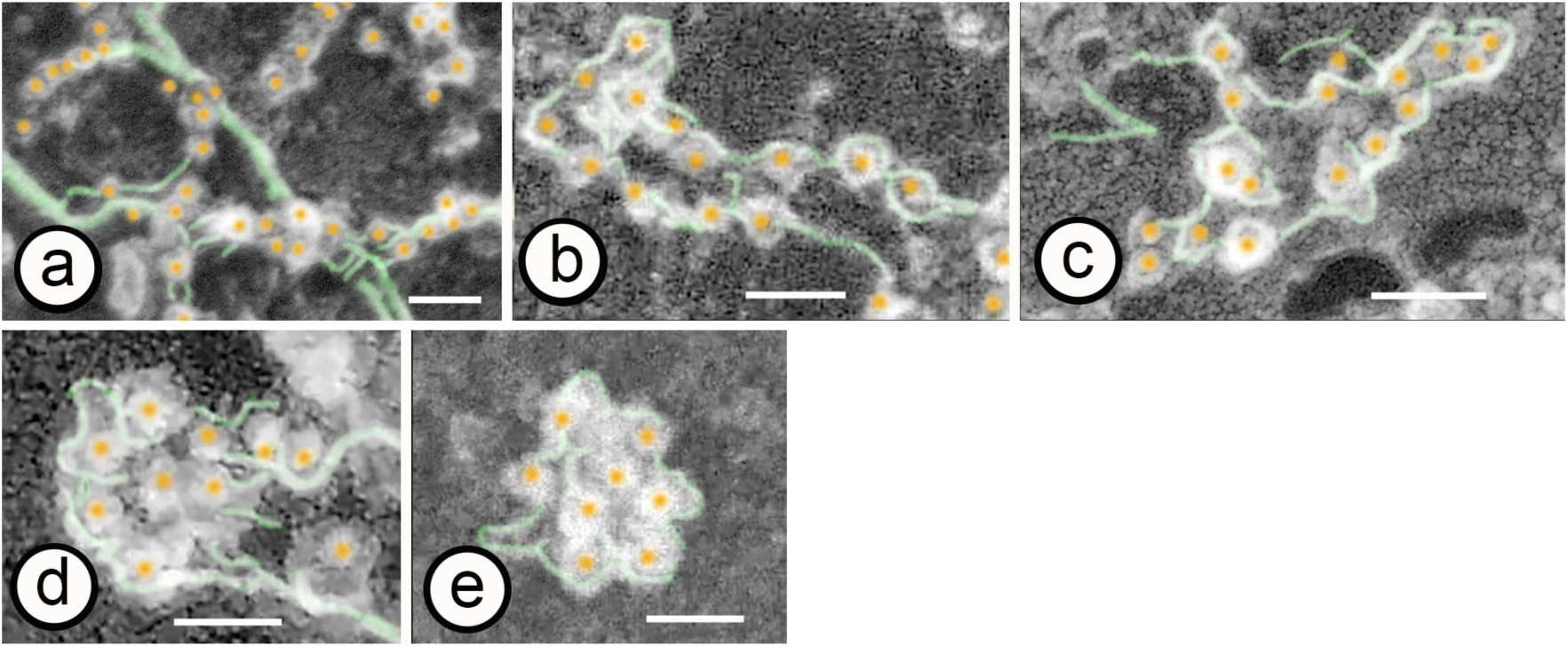
Gallery of immuno-FE electron micrographs showing various shapes of vRNP aggregates connected to each other by actin filaments, presumably indicating the process of the vRNP bundling/packaging (a to e). Scale bars: 50 nm

### The functionality of actin filaments in vRNP packaging

As noted above, the presence of myosin along actin filaments is suggestive of their active movement; to understand the vRNP packaging process more specifically, it is even more essential to know how and to what extent the actin filaments move. Atomic force microscopy (AFM) live cell imaging was therefore employed to visualise the movement of membrane cytoskeletal actin filaments just below the plasma membrane. Scanning the cantilever tip in close contact with the cell surface allows the detection of not only cell surface movements, such as lamellipodia, but also movements of actin filaments just below the plasma membrane ^35^. Fig. 5a shows a time-lapse image taken every 10 s, in which thin filaments just below the plasma membrane are moving randomly and rapidly (see also Extended Data Movie 1). These moving filaments are presumed to be actin filaments, judging from their thickness, movement, and sensitivity to blebbistatin, as discussed below. If this is the case, actin filaments bound to vRNPs observed in immuno-FE should also be actively moving in this way. Such active movement of actin filaments would facilitate mutual contacts between vRNPs bound to them, thereby permitting the sorting and assembly of eight distinct vRNPs. The presence of myosin along actin filaments (Fig. 3b) strongly suggests that movement of actin filaments is induced by interaction with myosin. Indeed, blocking myosin activity by blebbistatin reduced the movement of thin filaments (Fig. 5d, Extended Data Movie 2) and prevented proper assembly of vRNPs, often resulting in the appearance of large aggregates (compare Fig. 6a and Fig. 6b). Furthermore, morphological changes in vRNPs aggregates induced by blebbistatin treatment were statistically compared with those in cells cultured without blebbistatin (Fig.7). The distribution density of aggregates containing four or more anti-vRNP antibody labels (immuno-gold labels) on the cytoplasmic side surface of the plasma membrane was 25±5 aggregates / μm^2^ (mean value ± confidence limit (95%)) in the absence of blebbistatin, compared with 8±2 aggregates / μm^2^ (mean value ± confidence limit (95%)) in the presence of blebbistatin (Fig.7a). In addition, the number of aggregates (frequency of appearance) by size (number of antibody labels in each aggregate) is shown as a histogram (Fig. 7b). In the absence of blebbistatin, aggregates containing 7 to 9 antibody labels were the most abundant, forming a single sharp peak on the graph. However, in the presence of blebbistatin, no graphically significant peaks were formed, although the number of aggregates containing 10-12 antibody labels was slightly higher. Overall, aggregate formation of vRNPs in blebbistatin-treated cells shifted to a larger size. These results suggest a substantial involvement of actomyosin in vRNPs packaging. The significant differences were also found in the total number of anti-vRNP antibody labels distributed on the cytoplasmic side surface of the plasma membrane. In absence of blebbistatin, the number of the antibody labels per unit area (distribution density of antibody labels) was 231±28 labels /μm^2^ (mean value ±confidence limit (95%)), whereas in the presence of blebbistatin the distribution density of antibody labels was 149±28 labels /μm^2^ (mean value±confidence limit (95%))(Fig. 7a). In other words, inhibition of myosin activity reduces the total number of vRNPs reaching the cytoplasmic side surface of the plasma membrane. The actomyosin system may also be involved somehow in the transport of progeny vRNPs to the plasma membrane.

**Fig. 5.**
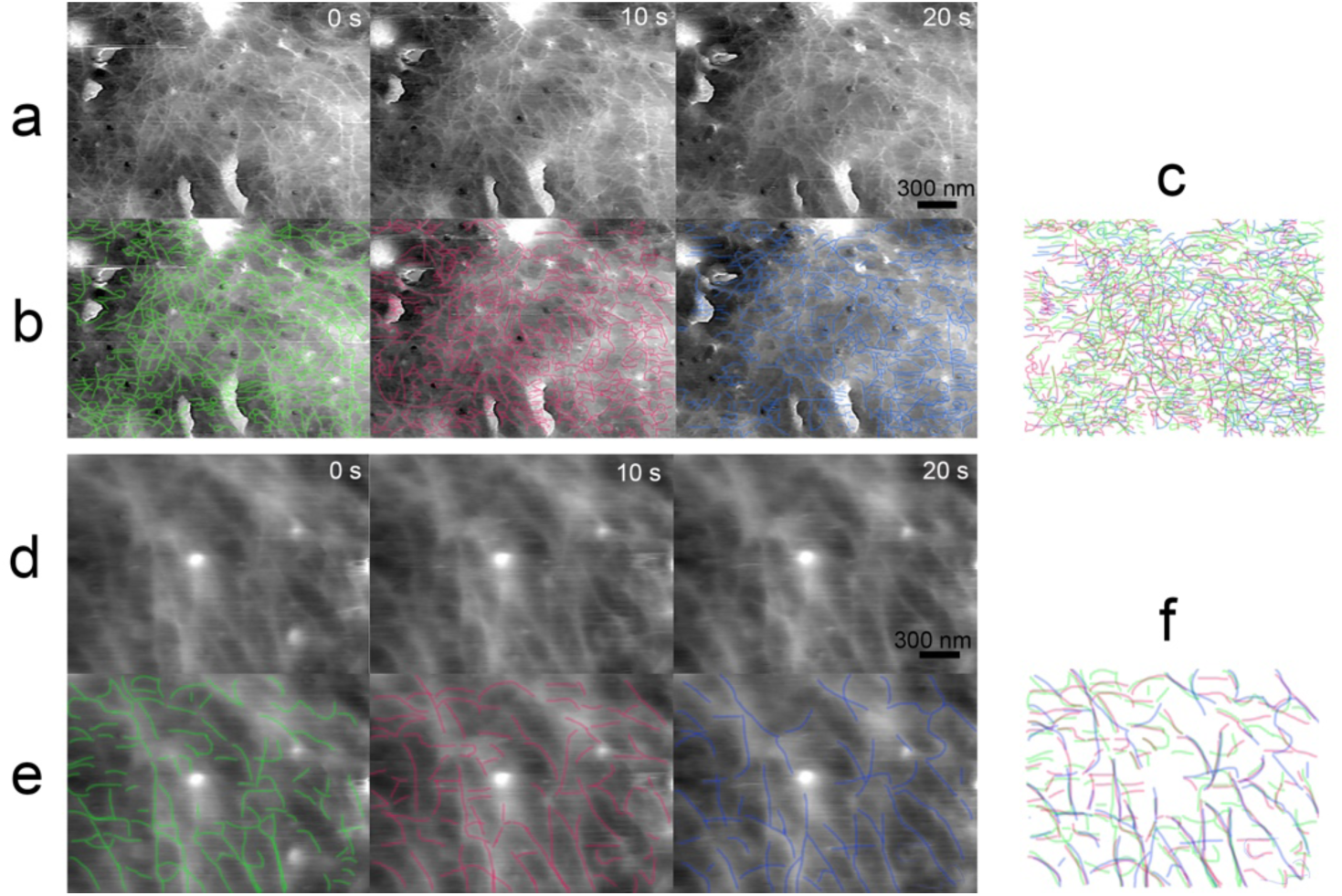
High-speed AFM time-laps images of IAV-infected cell surfaces showing the movement of thin filaments just below the plasma membrane in the absence (**a,b,c**) or the presence (**d,e,f)** of blebbistatin. The movement of thin filaments can be clearly detected by placing the cantilever in close contact with the cell surface. **a**: Time-lapse image at 10 s intervals in the absence of blebbistatin. **b:** The time-lapse image is the same as row a, but the thin filaments in each frame are traced in red, green, and blue respectively. **c:** The superimposed view of each colour trace line image in row b. The colours of the lines tracing thin filaments hardly overlap, so the number of lines traced appears to be increasing. This indicates that the thin filaments are actively moving. Compare with f below. **d**: Time-lapse image at 10 s intervals in the presence of blebbistatin. **e:** The time-lapse image is the same as row d, but the thin filaments in each frame are traced in red, green, and blue respectively. **f:** The superimposed view of each colour trace line image in row e. Each colour line image overlaps, indicating that the thin filaments have hardly moved. Scale bar: 300 nm

**Fig. 6.**
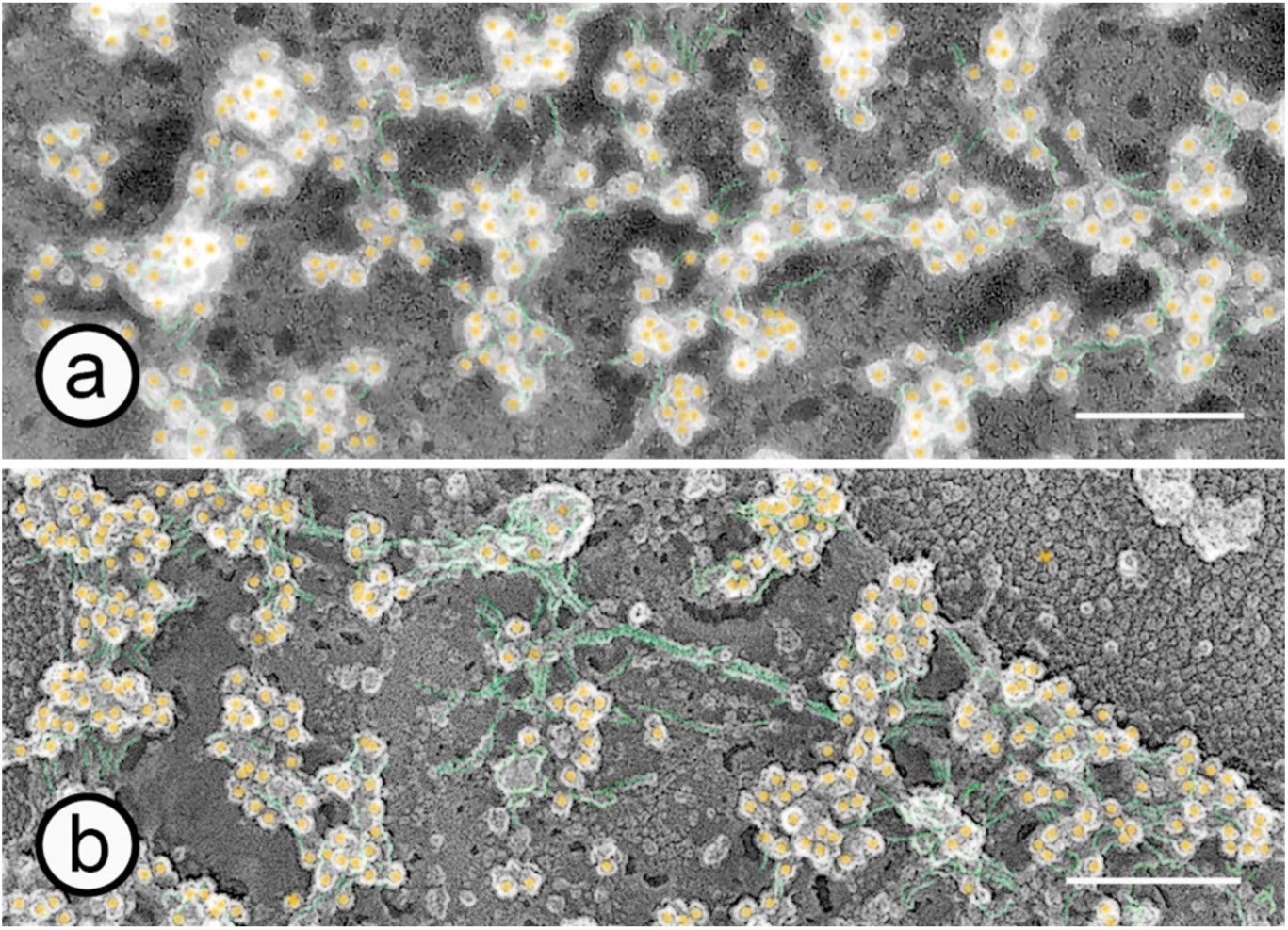
**a:** High magnification image of vRNP aggregates on the cytoplasmic side surface of the plasma membrane in cells infected with IAV in absence of blebbistatin. **b:** High magnification image of vRNP aggregates on the cytoplasmic side surface of the plasma membrane in cells infected with IAV in the presence of blebbistatin. Scale bars: 200 nm.

**Fig. 7.**
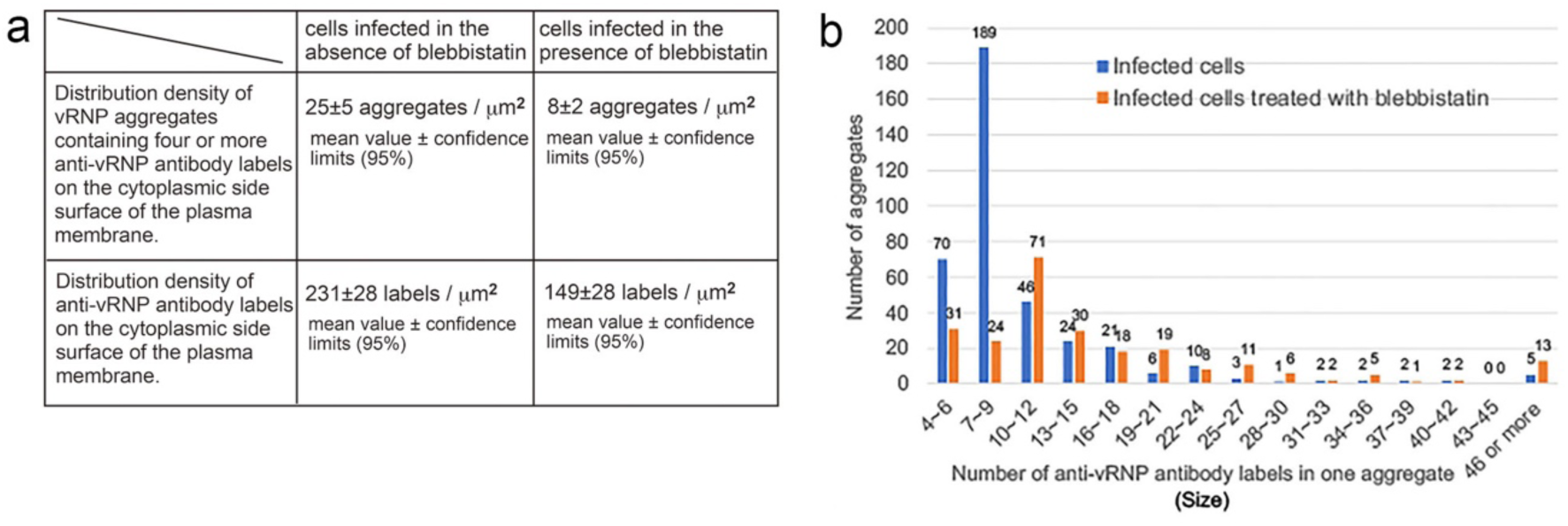
Statistical analysis of anti-vRNP antibody labels distributed and assembled on the cytoplasmic side surface of the plasma membrane. **a:** The table shows the distribution density of aggregates containing four or more vRNP antibody labels and of anti-vRNP antibody labels on the cytoplasmic side surface of the plasma membrane in the presence or absence of blebbistatin, respectively. In the presence of blebbistatin, not only the number of vRNP aggregates but also the total number of vRNPs that reach the plasma membrane is decreased. **b:** Histogram of aggregates by size. In the absence of blebbistatin, aggregates containing 7-9 labels are the most often detected, forming a sharp peak on the graph (blue bars). However, in the presence of blebbistatin, no significant peaks are formed on the graph (orange bars).

## Discussion

IAV is an enveloped, negative-strand RNA virus with a segmented genome that undergoes selective packaging, and therefore distinct vRNPs must interact specifically with each other to incorporate only one copy of each segment into a multi-segmented complex ^36–38^. If so, where and how are distinct vRNPs sorted and packaged into the core of the progeny virion? The present immuno-FE experiments of cells infected with IAV revealed that a large number of progeny vRNPs were accumulated on the cytoplasmic side surface of the plasma membrane, where they associated with actin filaments and were eventually packaged into circular aggregates through bundling of various shapes. It is therefore reasonable to assume that the cytoplasmic side surface of the plasma membrane is the assembly site where eight distinct vRNPs are sorted, packaged into the core of the progeny virion. Although the biochemical affinity of actin filaments to NP wrapping vRNA has been reported ^22,23^, the binding of actin filaments to vRNPs has not yet been visualised. This study was the first to visualise the association of vRNPs with actin filaments by using immuno-FE combined with unroofing technique. Even if this finding is the binding of the viral genome to actin filaments, it is of great interest from a cell biological point of view, as it contains the potential to be generalised to the affinity between the genome and actin filaments in normal cell nuclei. In the present experiments, vRNPs attached exclusively to actin filaments in contact with the cytoplasmic side surface of the plasma membrane but rarely attached to stress fibres, despite being composed of actin filaments. The reasons for this are currently unknown. It is also unclear how vRNPs bound to actin filaments are packaged into aggregates consisting of eight distinct vRNPs. Electron micrographs are still images and therefore unable to directly show the temporal processes of packaging. However, unless packaging process of vRNPs is synchronised, the different shapes of aggregates observed should reflect different stages of the packaging process. Figure 4 shows immuno-FE electron micrographs of various shapes of vRNP aggregates identified by anti-vRNP antibody labelling, pasted in sequence from linear arrays to tightly packed circular aggregates. Careful observation of many aggregate images allows us to infer the vRNP packaging process, which may proceed in the following order. i) vRNPs transported to the cell cortical area attach to actin bundles or actin filaments in contact with the cytoplasmic side surface of the plasma membrane. Actin bundles have been reported to dissociate laterally into single (thin) actin filaments by polyanions such as nucleotide phosphates or acidic amino acid oligomers^39^. As vRNPs are complexes of vRNAs (polynucleotides) and the NPs wrapping part of them, they are capable of working as polyanions that, when attached to the actin bundle, dissociate it laterally. ii) The transition from actin bundles to thin filaments makes it easier to move and reshape with the assistance of actin-binding proteins such as myosin, profilin and cofilin. Indeed, AFM live cell imaging showed active movement of actin filaments just below the plasma membrane. iii) Such movement may shuffle vRNPs on actin filaments, facilitating recognition by mutual contact. Eight distinct vRNPs are therefore presumed to be sorted and assembled in a manner dependent on the movement of actin filaments, such as elongation, bending and cleavage. iv) The assembly of eight distinct vRNPs results in more tightly bundled aggregates. v) Actin filaments between aggregates eventually disappear and each aggregate becomes independent. The process of packaging eight distinct vRNPs through these stages of bundling is illustrated as a hypothesis in Fig. 8. Whether this hypothesis is correct or not is currently uncertain, but it provides a good explanation for the existence of various aggregates of vRNPs bound to actin filaments. In AFM live cell imaging, inhibition of myosin activity by blebbistatin markedly prevented the movement of thin filaments (compare Figs 5a and 5b). Given that the AFM cantilever is only able to trace vertical irregularities of approximately 100 nm, it is strongly suggested that these moving thin filaments detected by AFM are actin filaments just below the plasma membrane, and that their movement is caused by the interaction between actin filaments and myosin. Inhibition of myosin activity also affected vRNP assembly, but less evident in comparison with Figs 6a and 6b. However, the effect of blebbistatin became clearer when the antibody-labeled vRNPs were counted and statistically processed (Fig. 7). Thus, myosin-induced movement of actin filaments appears to assist in the precise bundling eight distinct vRNPs. Furthermore, the total number of vRNPs arriving at the cytoplasmic side surface of the plasma membrane was also found to be significantly reduced in the presence of blebbistatin. This suggests that actomyosin motility is involved not only in vRNP packaging but also in transport. However, the transport pathway of progeny vRNPs from the nucleus to the cytoplasmic side surface of the plasma membrane is still unknown. In this study, antibody labelling of vRNPs was found in membrane-like structures of various sizes and shapes in the cytoplasm, but it is unclear whether these structures correspond to liquid organelles known as viral inclusions, as reported previously^16^. Although it is said that progeny vRNPs are transported on microtubules with the endoplasmic reticulum by cytoplasmic dynein, which is regulated by Rab11(small GTP binding protein)^16–20^. However, dynein carries cargo towards the minus end of microtubules (i.e. towards the centriole), the direction opposite to the plasma membrane^40^. Hence, transport towards the plasma membrane should be carried out by kinesins on microtubules or by myosin V on actin filaments^40^. However, transport by such kinesins and myosin is unlikely to be regulated by Rab11. Both intracellularly synthesised progeny vRNPs and progenitor vRNPs released from the newly entered viruses are mixed in the cell infected with IAV. Therefore, there are two directions of vRNPs trafficking; one towards the plasma membrane and the other towards the nucleus. The number of progeny vRNPs transported to the plasma membrane should increase rapidly with time after infection, but unfortunately our antibody is unable to distinguish progeny vRNPs from progenitor vRNPs. Rab11 should be always activated in infected cells because cytoplasmic dynein is also used to transport other cargo along microtubules in the direction towards the nucleus. In any case, further rigorous studies are required to elucidate the intracellular trafficking of vRNPs.

**Fig. 8.**
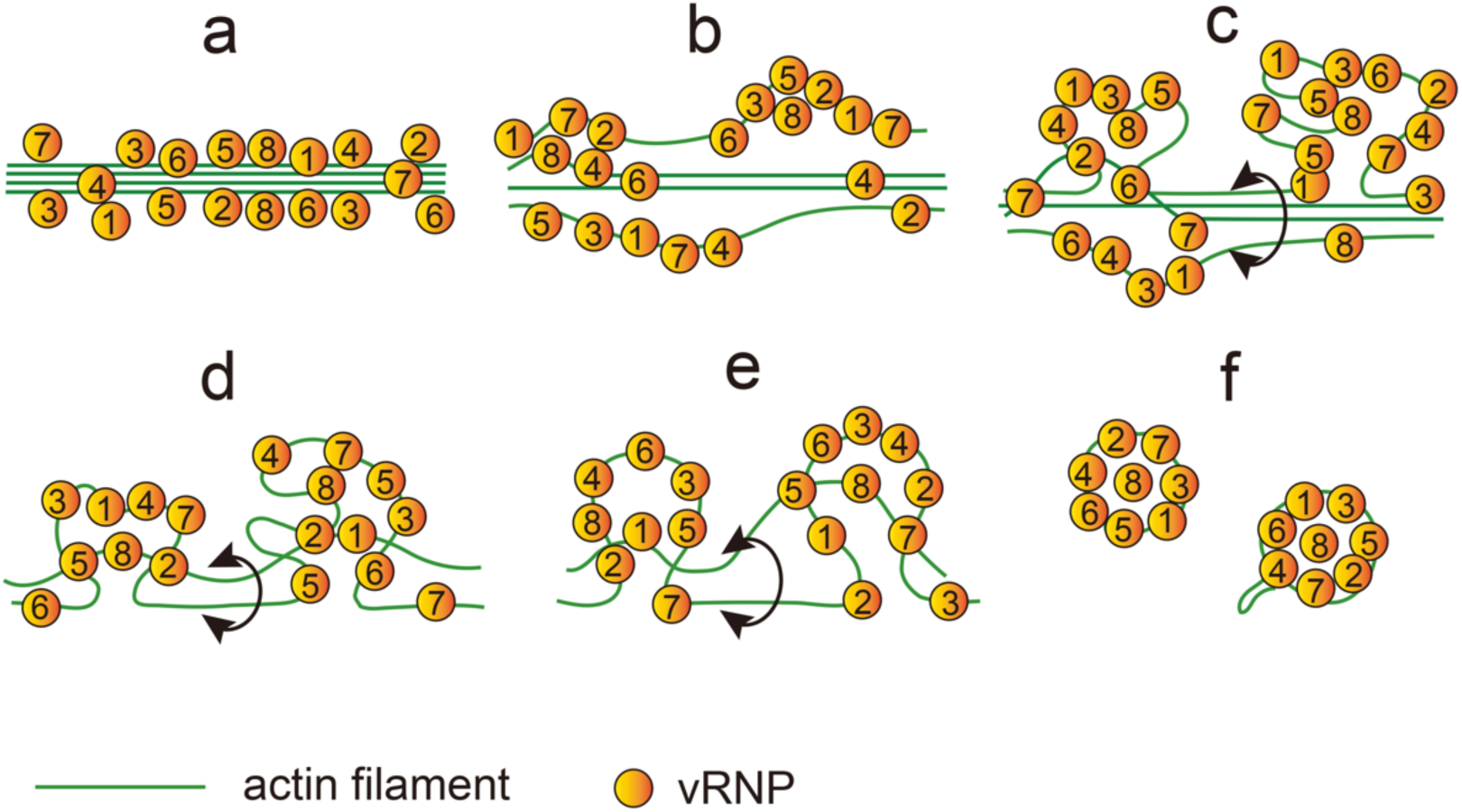
Hypothetical diagram showing the process in which eight distinct genomic vRNPs are packaged by actin filaments. **a:** Alignment of vRNPs on actin bundles appears to be the first step. **b:** Polyanions, such as polynucleotides, are thought to laterally separate thick actin bundles to thin actin filaments ^39^, resulting in the translocation of vRNPs to thin actin filaments. **c, d:** vRNPs aligned on thin actin filaments gradually aggregate by myosin-induced movement of the actin filaments. The mutual contact of vRNPs with each other facilitate selective packaging through vRNP-vRNP interaction. **e:** The parts of the aggregate composed of distinct vRNPs begin to tighten, and overlapping vRNP segments loosen and are excluded from the aggregate. This process is expected to be repeated until eight distinct vRNPs have aggregated. **f:** Once eight distinct vRNPs have assembled, they appear to be more tightly packed with the assistance of actin filaments. Eventually, actin filaments between aggregates (encircled by double head curved arrows) disappear. The number given to the vRNPs (1-8) is arbitrary.

Actin and myosin each have several isoforms with different functions and localisation. The antibody used to investigate actin localisation in this study was an anti-cytoskeletal γ-actin antibody, but actin is a very conserved molecule, and it is difficult to distinguish isoforms with antibodies. The same is true for myosin. Anti-myosin II antibody used in this study is also thought to label multiple isoforms. Therefore, in this paper, we simply used the terms actin and myosin without selecting the names of the antigenic isoforms.

Since progeny virions bud from the apical surface of infected cells in the respiratory epithelium, the cytoplasmic side surface of the dorsal plasma membrane is expected to be different from that of the ventral plasma membrane. Unlike cells in tissue, however, cells in culture are thought to have already lost their intracellular functional and structural polarity. Hence, we observed whether the structure of the cytoplasmic surface of the dorsal plasma membrane is different from that of the ventral plasma membrane, using an adhesive unroofing technique in which the dorsal plasma membrane is pulled off with an adhesive coverslip (Extended data Fig. 2c). Consequently, both dorsal and ventral plasma membranes were structurally equivalent. Like the ventral plasma membrane, vRNPs linked by actin filaments formed aggregates on the cytoplasmic side surface of the dorsal plasma membrane (Extended data Fig.3). Not surprisingly, in cultured cells, progeny virions also budded from the ventral plasma membrane to the substrate side. (Extended data Fig. 4). These facts suggest that some of the spatial polarities inherent to cells in tissue are lost in cells cultured.

This study used advanced immuno-FE and high-speed AFM to reveal the interplay between membrane cytoskeletal actin filaments and genomic vRNPs. Further deepening of our understanding of the mechanisms involved in IAV genome packaging in host cells at the molecular level will assist the development of antiviral strategies that complement vaccines.

## Methods

### Cells and viruses

For immuno-FE, A549 cells derived from human lung epithelial cells (ATCC, VI, USA) were cultivated in Ham’s F-12K medium (Wako, Osaka, Japan) supplemented with 10% foetal calf serum (FCS) and penicillin‒streptomycin (100 unit/ml and 100 µg/ml, respectively). Influenza virus strain A/WSN/1933 (H1N1; WSN) was generated in 293T human embryonic kidney cells by employing a plasmid-based reverse genetics system as described previously ^41^. The 293T cells were maintained in Dulbecco’s modified Eagle’s medium (DMEM, Sigma‒Aldrich, USA) with 10% FCS and penicillin‒streptomycin. Propagation and titration of the WSN virus were conducted in Madin-Darby canine kidney (MDCK) cells. The MDCK cells were maintained in Eagle’s minimum essential medium (MEM) containing 5% new born calf serum (NCS) and penicillin‒streptomycin, but during virus propagation and titration, MEM supplemented with 0.2% bovine serum albumin (BSA) was used instead of serum-supplemented medium. All cells were cultured at 37 °C under 5% CO_2_. For AFM live-cell imaging, we used a replication-incompetent PB2-knockout influenza A/Puerto Rico/8/34 (H1N1; PR8) virus for biosafety reasons. This virus also possessed a green fluorescent protein (GFP) reporter gene (PB2-KO/GFP virus) and was generated by using reverse genetics^41^. The PB2-KO/GFP virus was propagated and titrated in AX4/PB2 cells, an MDCK cell derivative overexpressing human 2,6-sialyltransferase and PB2 of the PR8 virus ^42^. AX4/PB2 cells can complement viral growth and were therefore also used for observation of virus budding. The AX4/PB2 cells were maintained in MEM containing 5% NCS and penicillin‒streptomycin.

### Immuno-freeze etching replica (Immuno-FE) of cryo-sections

Immuno-FE of a cryo-section was performed to examine the cytoplasmic distribution of vRNP in cells infected with IAV. A549 cells were cultured in a plastic dish (ø 9 cm). Cells were grown to ∼80% confluence in the dish and then infected with WSN virus at a multiplicity of infection (MOI) of 10 for 16 h. Cultured cells were washed once with phosphate-buffered saline (PBS) and immediately fixed with 2% glutaraldehyde in the KHMgE buffer (70 mM KCl, 5 mM MgCl_2_, 3 mM EGTA in 30 mM HEPES buffer, adjusted to pH 7.4 with KOH) for 20 min. Cells were harvested with a cell scraper and washed with PBS. The resulting pellet was soaked in 2.3 M sucrose for at least 3 h to infiltrate the pellets. The sample (pellet) was then mounted on a circular sample carrier, a consumable for the Leica EM FC 7 cryo-microtome (Leica Microsystems, Wetzlar, Germany), and frozen in liquid nitrogen. The experimental process up to preparing the frozen sections was carried out according to the Tokuyasu method ^28, 29^. Frozen sections approximately 120-nm thick were collected on 2.5 x 2.5 mm glass coverslips (#1 thickness, Matsunami Glass Industry, Ltd., Osaka, Japan) (Extended data Fig.2a), washed three times with PBS after bringing to room temperature, and immersed in 4% BSA for blocking. The sections were incubated with an anti-vRNP polyclonal antibody (available in the Kawaoka Laboratory) overnight at 4 °C. The polyclonal antibodies were diluted with PBS containing 1% BSA at a ratio of 1:200. After labelling with the primary antibody, the sections were washed with fresh PBS three times for 5 min each and incubated with 10 nm colloidal gold-conjugated goat anti-rabbit IgG for 2 h (EM. GARG 10, British BioCell International, Scotland, UK). The antibody-labelled cells were washed three times with PBS for 5 min each and fixed again with 2% glutaraldehyde for at least 10 min. Subsequently, the cells were frozen rapidly by plunging them onto a copper block cooled with liquid helium immediately after being washed twice with distilled water ^30^. Immuno-labelled frozen sections were brought into the FR-9000 freeze-etching equipment (Hitachi High-Tech, Tokyo, Japan) and deep-etched (slight freeze-drying) at −90℃ for approximately 10 min, after which platinum and carbon were evaporated to produce a replica of the surface structure.

### Immuno-FE of unroofed cells

A549 cells were cultured on 2.5 x 2.5 mm coverslips as described above. The cells were grown to approximately 80% confluence and then infected with WSN virus at an MOI of 10 for 16 h. Some infected cells were cultured in medium containing blebbistatin (Sigma-Aldrich, St Louis, MO, USA) dissolved in 1% dimethyl sulphoxide (DMSO). Blebbistatin was added to the medium to a final concentration of 50 µM 3 hours before unroofing. Infected cells and non-infected cells (control or mock-infected) on cover slips were unroofed by sonication to observe the cytoplasmic side surface of the ventral plasma membrane (Sonication unroofing, Extended data Fig. 2 b), as described previously ^31, 32^. Prior to sonication, cell adhesion to the coverslip was enhanced by treatment with poly-L-lysine (Mw 30,000–50,000; Sigma‒Aldrich, USA; 0.5 mg/ml in calcium-free Ringer’s solution) for 5 seconds, and the cells were washed in calcium-free Ringer’s solution for a few seconds. Subsequently, the cells were further washed in KHMgE buffer for a few seconds, and then exposed for 5–10 seconds to fine air bubbles (cavitation) generated by weak sonication (0.3–0.5 W, 27 kHz) in the KHMgE buffer containing protease inhibitor (cOmplete Sigma‒Aldrich, USA). Cavitation ruptures the dorsal plasma membrane and removes cytoplasm, so that the cytoplasmic side surface of the ventral plasma membrane is exposed.

Another unroofing method (adhesion unroofing)^32^ was employed to observe the cytoplasmic side surface of the dorsal plasma membrane (Extended Data Fig. 2c). In the adhesion unroofing, the dorsal plasma membrane is peeled off using the adhesive properties of the coverslips treated with Alcian blue. For making adhesive coverslips, coverslips (2.5 mm x 2.5 mm), hydrophilised by ion cleaning with plasma sputtering, were immersed in a 1% Alcian blue solution and treated for approximately 1 minute. The Alcian blue solution on the coverslips was then washed off with distilled water and dried before use^32^. Adhesive coverslips are placed in close contact with the apical cell surface. Excess amount of buffer around the coverslip was absorbed with filter papers; after standing for 30 s, fixative (approx. 250 µL) (2% glutaraldehyde in KHMgE buffer) was applied. The dorsal plasma membrane is peeled off onto the coverslip by buoyancy force from the fixative solution or by gently pulling up with tweezers. This allows the cytoplasmic side surface of the dorsal plasma membrane to be exposed for observation, but the yield is low because it depends on the adhesiveness of the coverslip surface. Regardless of which method was used for unroofing, fixation and immunolabeling were carried out in the same way. Unroofed cells were fixed in 2% glutaraldehyde in the KHMgE buffer for at least 20 min. Experiments from unroofing to fixation were carried out in a safety cabinet.

For immunolabelling, unroofed and fixed cells on coverslips were washed twice with PBS and soaked in 4% BSA in PBS for 10 min to block nonspecific binding. The cells were then incubated with the anti-vRNP polyclonal antibody (produced in Kawaoka lab.), anti-cytoskeletal γ actin polyclonal antibody (Novus Biological LLC, CO USA) or anti-myosin II non-muscle type polyclonal antibody (Sigma-Aldrich MO USA) overnight at 4 °C. The antibody was diluted with PBS containing 1% BSA at ratios of 1:200. After labelling with the primary antibody, the cells were washed with fresh PBS three times for 5 min each and then incubated with 10 nm gold-conjugated goat anti-rabbit IgG (British BioCell International, UK) for 2 h. As a control for immunolabelling, nonimmune rabbit serum (Invitrogen MA USA) was used instead of primary antibody. Freeze-etched replicas were obtained as described above using the FR-9000 freeze-etching machine. Photoshop, Fiji, and Excel were used to quantify immuno-FE micrographs.

### AFM live cell imaging

AX4/PB2 cells (MDCK cells genetically modified to express PB2 as described above) were cultured in 500-μL droplets of MEM on glass slides edged with hydrophobic ink (TF0215; Matsunami Glass Industry, Ltd. Osaka Japan). After the cells were grown to approximately 70% confluence, they were infected with PB2-KO/GFP virus at MOI 5 and incubated for a further 10 hours.

To examine actin filament motility after blocking myosin activity, blebbistatin was dissolved in 1% DEMSO and added to the culture medium to a final concentration of 50 µM. Cells were incubated for further 3 hours and then observed. The cells on the glass slides were imaged in a tip-scan-type high-speed AFM imaging system that was an improved version of a previously developed AFM (BIXAM; Olympus Corporation, Tokyo, Japan) ^43^. AFM was performed in the phase-modulation mode using small and soft cantilevers (2-μm wide, 9-μm long, and 0.1-μm thick) with a spring constant of 0.1 N/m (USC-F0.8-k0.1-T12-10: NanoWorld AG, Neuchatel, Switzerland). The diameter of the scanning tip was approximately 8 nm. The force against samples was estimated to be 50 pN. Time-lapse image series were acquired by scanning a 6.0 μm (x-axis) × 4.5 μm (y-axis) area of the cell surface at a scanning rate of 0.1 frames per second (fps). The scanning resolution was 320 pixels (x-axis) and 240 lines (y-axis).

### Western blot analysis

Viral or A549 cell extracts were prepared and mixed with 4x SDS sample buffer (Bio-Rad, CA, USA) supplemented with 2 mM DTT. The same amounts of viral and A549 cell extracts were analysed by a general protocol with slight modifications as reported previously^44^. Then, the gels were transferred onto PVDF membranes using the semidry method (BIO CRAFT, Tokyo, Japan) in different blotting buffers [(A) 300 mM Tris, 20% Et-OH, 0.02% SDS; (B) 25 mM Tris, 20% Et-OH, 0.02% SDS; and (C) 25 mM Tris, 20% Et-OH, 40 mM 6-aminohexanoic acid, 0.02% SDS) for 2 h at 50 mA (constant current)]. After blocking with skimmed milk in 1 x RBS for 1 h, a primary antibody (anti-cytoskeletal γ-actin polyclonal antibody NB600-533 (NOVUS Biological CO, USA), anti-myosin IIa nonmuscle polyclonal antibody M8064 (Sigma, MO, USA) anti-cofilin antibody ab42824 (Abcam, MA, USA), anti-M1 monoclonal antibody HB64 or anti-NP antibody HB65 (ATCC, VI, USA) was used. Finally, the signals were detected with Immobilon® Forte Western HRP Substrate (Merck, Darmstadt, Germany). The chemiluminescent signals were detected with a LuminoGraph II EM (ATTO, Tokyo, Japan).

## Acknowledgements

This work was supported, in part, by the Development of Advanced Measurement and Analysis Systems (AMED-SENTAN) (#18hm0102003s0107 to J.U.) program funded by the Japan Agency for Medical Research and Development and by Grants-in-Aid for Scientific Research (#20K06586 to J.U. and #19K06583 to E.U.) from JSPS and a JBUK Award from JSPS London to Y.Y. We thank Atzin Bolanos-Ceron for HB64 and HB65 monoclonal antibody purification from hybridoma supernatants.

## Author contributions

All authors were involved in this study in their respective areas of expertise. H-I.W. prepared infected cell samples for immuno-freeze-etching EM and AFM. J.U. performed cryo-sectioning and unroofing of infected cells for immuno-FE, AFM and morphological analysis of immuno-FE. E.U. performed AFM live-cell imaging with A.N., Y.Y. and H-I.W. Y.M. performed western blot analysis. The manuscript was mainly written by J.U., Y.Y. and Y.K. with contributions from all authors. Conceptualization of the study: J.U., Y.K., H-I.W., and Y.Y.

## Additional information, Supplementary information

The online version contains methods and extended data.

## Data availability

All data are available from the corresponding authors on request.

## Extended data

**Extended data Fig. 1.**
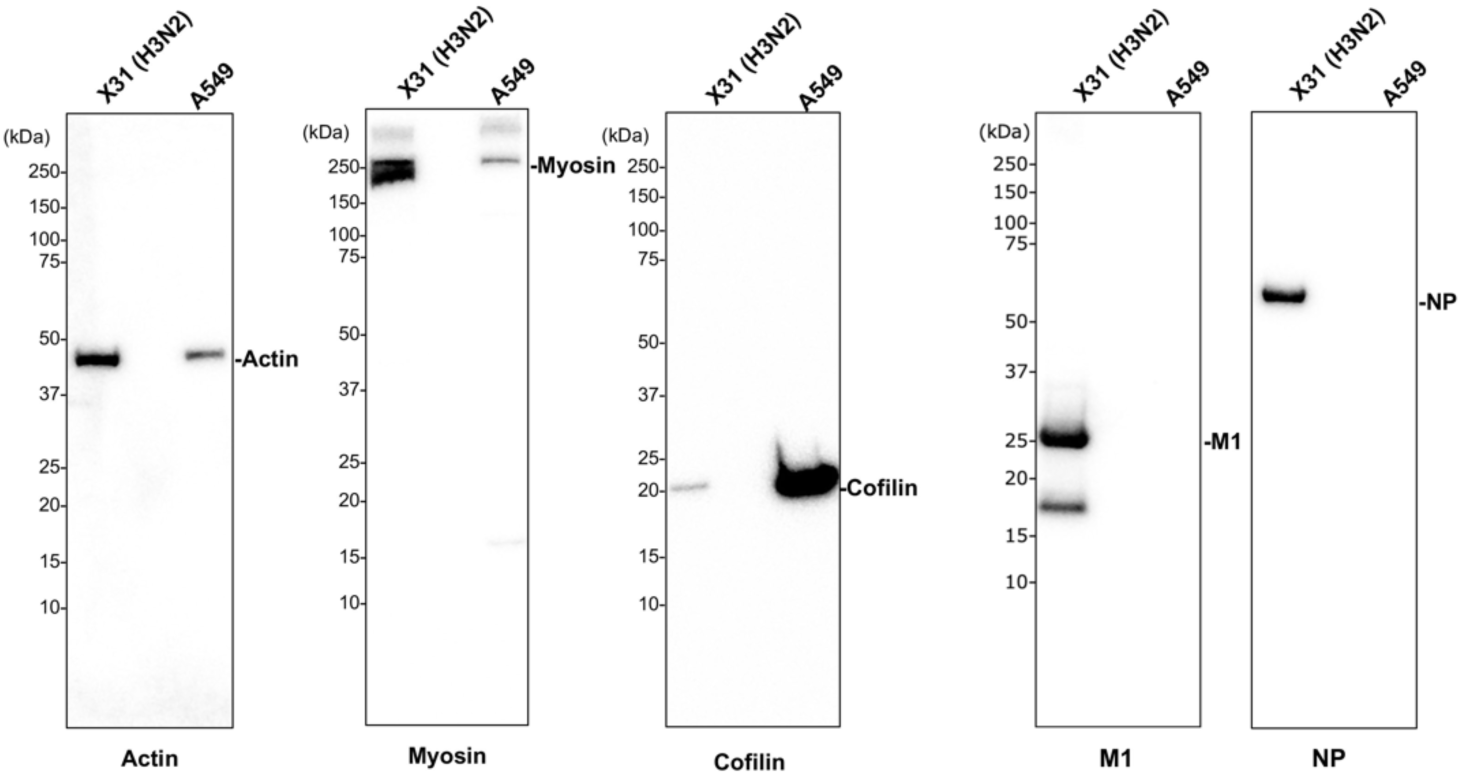
Western blot analysis showing the presence of actin, myosin, and cofilin in purified IAV (X31) virions and non-infected A549 cells, which were detected by using an anti-cytoskeletal (γ) actin polyclonal antibody, an anti-myosin II polyclonal antibody, and an anti-cofilin-1 polyclonal antibody, respectively. The same amount of sample was loaded in each lane. Actin and myosin were detected in both A549 uninfected cells and IAV. Cofilin was also present in the IAV, but in smaller quantities. The IAV-specific proteins M1 and NP were also examined by Western blotting as references. Neither protein was present in non-infected A549 cells.

**Extended data Fig. 2.**
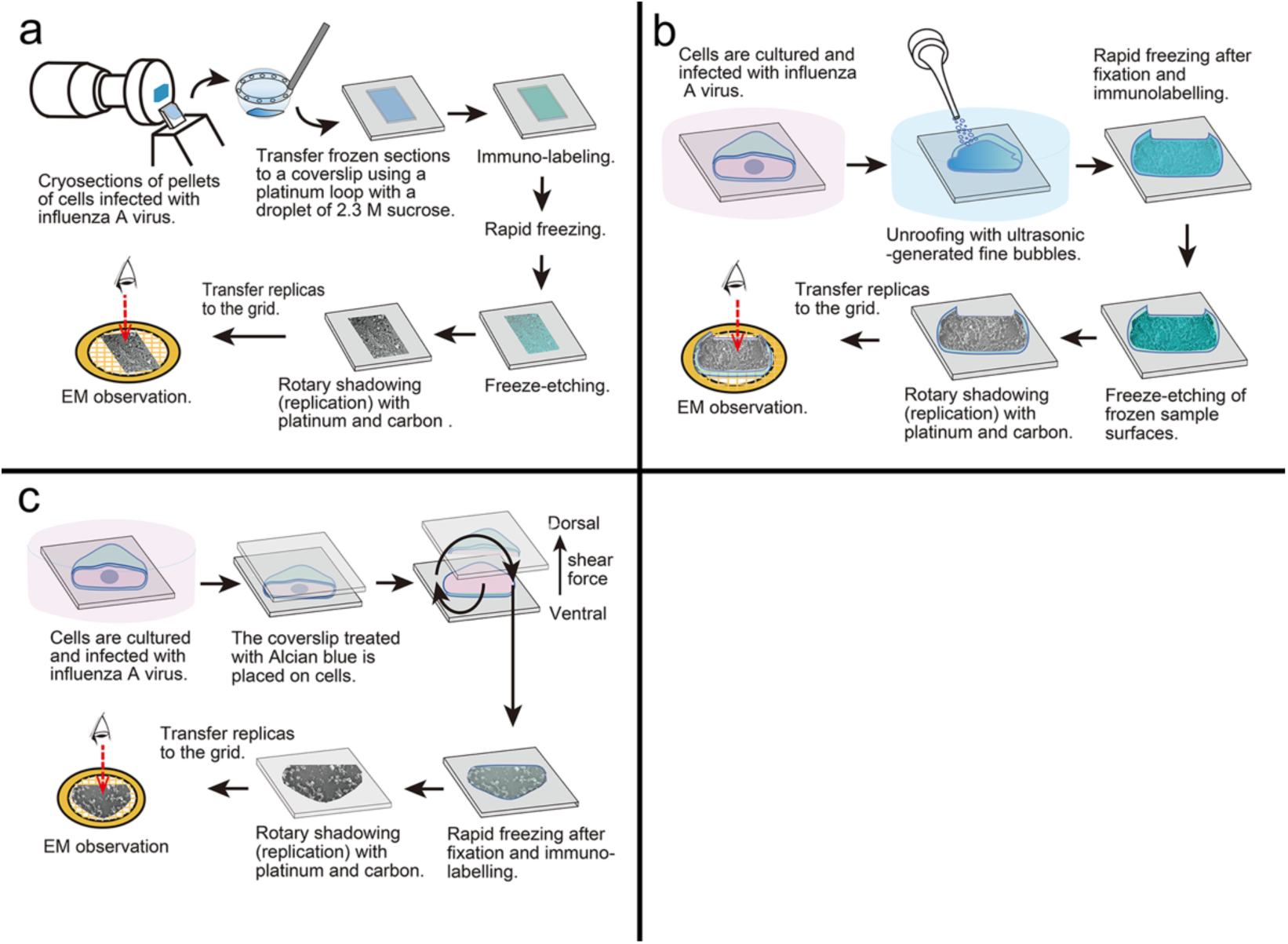
Illustration showing the procedure of cryo-section immuno-FE (a), sonication unroofing immuno-FE (b) and adhesion unroofing immuno-FE (c). **a:** Cryo-sections are transferred onto small coverslips and labelled with anti-vRNP antibody. The labelled sections are quickly frozen again and freeze-etched (freeze-dried). The surface structure of the section is replicated by evaporation of platinum and carbon. **b:** The dorsal plasma membrane and cytoplasm are removed with fine bubbles generated by sonication in the buffer. The remaining ventral plasma membrane is fixed and labelled with anti-vRNP antibody, anti-myosin antibody or anti-actin antibody. The samples are quickly frozen, freeze-etched, and replicated by evaporation of platinum and carbon. **c:** Adhesion unroofing is an alternative unroofing method in which an Alcian blue-treated sticky coverslip is pressed on cells and then lifted to peel off the dorsal cell membrane onto the coverslip. Hence, adhesion unroofing can harvest the dorsal plasma membrane and expose its cytoplasmic side surface. The exposed surface is then labelled with anti-vRNP antibodies and replicated in the same way as described above.

**Extended data Fig. 3.**
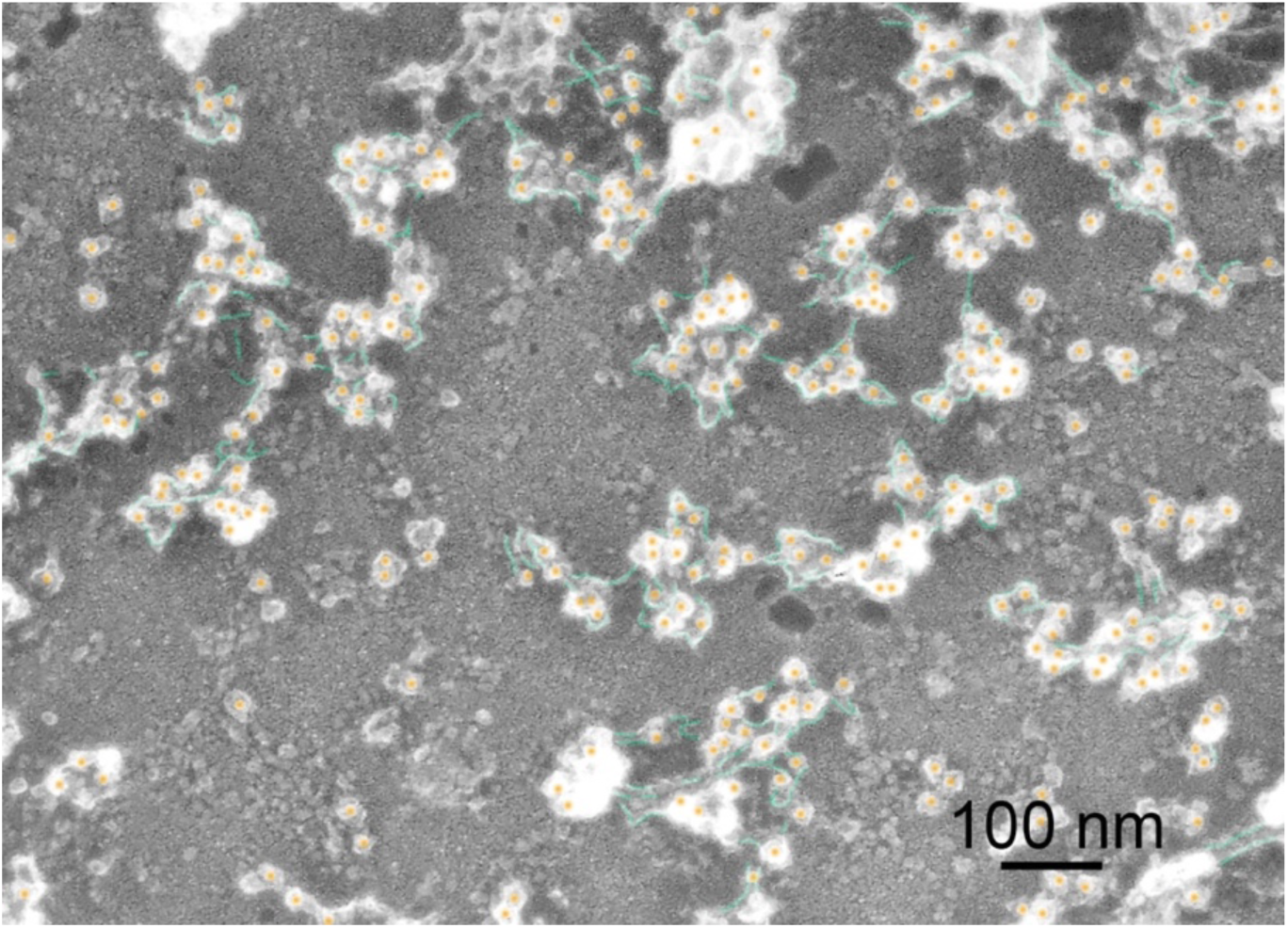
Immuno-FE micrograph showing the cytoplasmic side surface of dorsal plasma membrane of infected cells exposed by adhesion unroofing. Anti-vRNP antibody labels (i.e. vRNPs, coloured in orange) are aggregated linking with actin filaments (coloured in green). Such structural appearance is identical to that of the cytoplasmic side surface of the ventral plasma membrane.

**Extended data Fig. 4.**
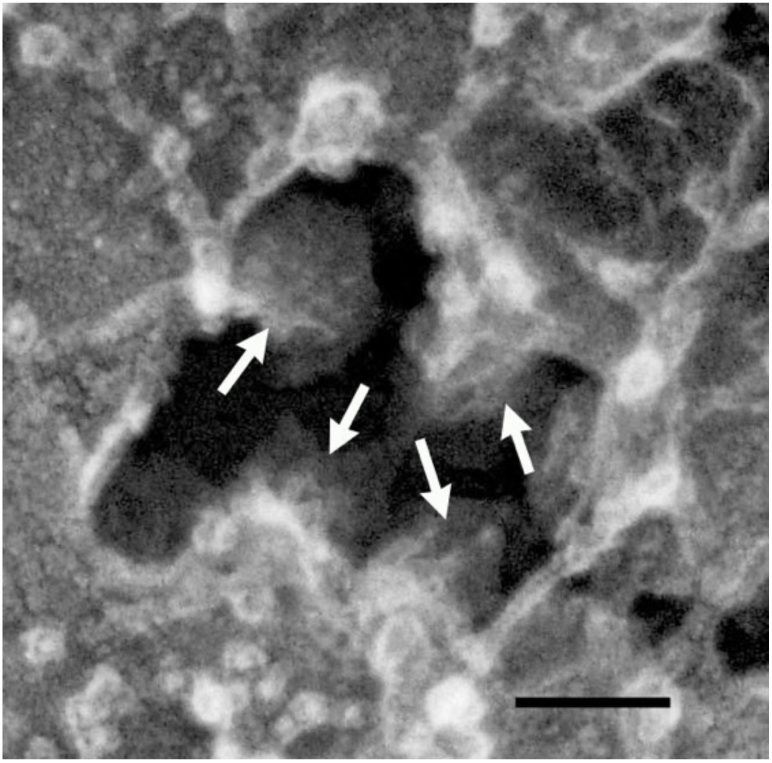
Freeze-etched electron micrograph showing progeny virions (arrows) budding from the ventral plasma membrane to the substrate side. The ventral plasma membrane is partially broken, from which the progeny virions can be seen budding out to the substrate side. Scale bar: 100 nm

Extended data video 1: AFM live-cell imaging showing rapid movement of actin filaments just below the plasma membrane.

Extended data video 2: AFM live-cell imaging showing prevention of motility of actin filaments just below the plasma membrane in the presence of blebbistatin.

